# Relationship between neural activity and neuronal cell fate in regenerating *Hydra* revealed by cell-type specific imaging

**DOI:** 10.1101/2023.03.19.533365

**Authors:** Alondra Escobar, Soonyoung Kim, Abby S. Primack, Guillaume Duret, Celina E. Juliano, Jacob T. Robinson

## Abstract

Understanding how neural circuits are regenerated following injury is a fundamental question in neuroscience. *Hydra* is a powerful model for studying this process because it has significant and reproducible regenerative abilities, a simple and transparent body that allows for whole nervous system imaging, and established methods for creating transgenics with cell-type-specific expression. In addition, cnidarians such as *Hydra* split from bilaterians (the group that encompasses most model organisms used in neuroscience) over 500 million years ago, so similarities with other models likely indicates deeply conserved biological processes. *Hydra* is a long-standing regeneration model and is an emerging model for neuroscience; however, relatively little is known regarding the restoration of neural activity and behavior following significant injury. In this study, we ask if regenerating neurons reach a terminal cell fate and then reform functional neural circuits, or if neural circuits regenerate first and then guide the constituent cells toward their terminal fate. To address this question, we developed a dual-expression transgenic *Hydra* line that expresses a cell-type-specific red fluorescent protein (tdTomato) in ec5 peduncle neurons, and a calcium indicator (GCaMP7s) in all neurons. This transgenic line allowed us to monitor neural activity while we simultaneously track the reappearance of terminally differentiated ec5 neurons as determined by the expression of tdTomato. Using SCAPE (Swept Confocally Aligned Planar Excitation) microscopy, we tracked both calcium activity and expression of tdTomato-positive neurons in 3D with single-cell resolution during regeneration of *Hydra’s* aboral end. We observed tdTomato expression in ec5 neurons approximately four hours before the neural activity begins to display synchronized patterns associated with a regenerated neural circuit. These data suggest that regenerating neurons undergo terminal differentiation prior to re-establishing their functional role in the nervous system. The combination of dynamic imaging of neural activity and gene expression during regeneration make *Hydra* a powerful model system for understanding the key molecular and functional processes involved in neuro-regeneration following injury.

## Introduction

Neural regeneration capacity is widely exemplified across animals. The extent of these regenerative capacities ranges from cellular regeneration to whole body reformation, with some of the most extreme examples of nervous system regeneration found in flatworms (e.g., *Schmidtea mediterranea)* [1,2], cnidarians (e.g., *Hydra vulgaris*) [3,4,5], the replacement of complete innervated limbs in salamanders (e.g., *Ambystoma mexicanum*) [6,7] or spinal cord injury recovery in zebrafish (e.g., *Danio rerio*) [8,9]. By contrast, mammals exhibit limited regenerative abilities, along with a complex immune response that slows neural regrowth [10]. Understanding the molecular and cellular mechanisms that drive nervous system regeneration in highly regenerative animals will likely inform the development of neural repair therapies for humans [11]. In particular, we need to understand how newly regenerated neurons rebuild functional neural circuits. The cnidarian polyp *Hydra* has a simple neural structure, extensive neuron regenerative capabilities [12,13], established genetic tools [14,15], and is an emerging neuroscience model [16]. These features make *Hydr*a an excellent model for interrogating the cellular dynamics of neural circuit functional regeneration.

*Hydra* has a simple radial body plan organized around a single oral-aboral axis of symmetry, with the hypostome and tentacles at the oral end (i.e., the head) and the peduncle and basal disk at the aboral end. The *Hydra* body is formed by two epithelial monolayers, the inner endoderm and outer ectoderm, separated by an extracellular matrix [17]. *Hydra* has two separate nerve nets, one embedded in each of the epithelial layers. Neurons run along the entire length of the body, with a higher neuron density in the hypostome (oral end) and the peduncle and basal disc (aboral end) [18,41]. Important groundwork has been done to identify neural circuits associated with specific behaviors in *Hydra* [19]. This aids our investigation of neural circuit regeneration because the resumption of normal behavior indicates when a neural circuit has functionally regenerated. Specifically, *Hydra* has a defined repertoire of behaviors associated with four major non overlapping neural circuits [28]. In this study we focus on longitudinal contractions, because they are the easiest movements to track and are known to correlate with the activity of the Contraction Burst (CB) circuit. The CB circuit involves neurons that run the length of the ectodermal epithelium, including a particularly prominent group of neurons located in the peduncle at the aboral end [19].

The transcriptional state of all eleven *Hydra* neuron subtypes has been profiled using single cell RNA-sequencing [1, 47]. Similar to findings in *C. elegans* [20], the *Hydra* neurons are best defined by unique combinatorial expression of specific genes, including transcription factors and neuropeptides [21,22]. Previous work suggests that the neuron subtypes that participate in the CB circuit are defined by combinatorial expression of various paralogs of the *hym176* neuropeptide gene (*hym176A-E*) [14,23]. One of these neuron subtypes is the “ec5” population, which is located in the peduncle and selectively expresses the neuropeptide Hym176C [24,25].

In addition, cnidarians such as *Hydra* split from bilaterians (the group that encompasses most model organisms used in neuroscience) over 500 million years ago, so similarities with other models likely indicates deeply conserved biological processes. *Hydra* is a long-standing regeneration model and is an emerging model for neuroscience thanks to recent studies that elucidate stem cell differentiation trajectories in *Hydra* cells, including all neuronal subtypes [26]. However, relatively little is known about the restoration of neural activity [27,28,29] and behavior following a significant injury. As part of the regeneration process of a neural circuit, two key events occur: terminal cell differentiation and the synchronization of cell activity which elucidates recovery behavior[]. In *Hydra*, the order of these key events during its neural regeneration process remained unknown. In this study, we ask if regenerating neurons reach a terminal cell fate and then reform functional neural circuits, or if neural circuits regenerate first and then guide the constituent cells toward their terminal fate.

In this study, we use the regulatory region of hym176c to drive ec5-specific nuclear expression of tdTomato along with GCaMP7s [30] in all neurons. This allowed us to perform cell-type specific 3D imaging of neural activity in both uninjured and regenerating *Hydra*. We first confirmed that ec5 neurons act in the CB circuit, and then observed the regeneration of the CB circuit by tracking the neural activity and reappearance of ec5 neurons after foot amputation. We found that ec5 neurons terminally differentiate before they synchronize with neighboring neurons. This suggests that positional cues, not neural activity cues play the dominant role in guiding neuronal cell fate in this circuit. This study provides the foundational tools and conceptual framework to better understand the molecular mechanisms that underlie the regeneration of functional neural circuits.

## Results and Discussion

### Ec5 neurons are a part of the Contraction Burst (CB) Circuit

To monitor the activity of the ec5 neurons both in uninjured *Hydra* and during regeneration, we used transgenic line *Tg(hym176c:tdTomato,tba1c:GCAMP7s)*^cj1-in^, which was created for this study, and in which tdTomato is specifically expressed in ec5 neurons (Fig. 1). To create this line we extracted the regulatory region of the *hym176c* gene, which is expressed specifically in ec5 neurons (Figure 1h) and used this to drive the expression of nuclear tdTomato. To be able to track ec5 expression in the context of the activity of the entire nervous system, we included a second expression cassette in the construct using GCaMP7s under the control of pan-neuronal promoter *tba1c (*identified in [47]). After obtaining a founding polyp with integration of the transgene in the interstitial lineage, which includes all neurons, we propagated the line through asexual reproduction, which gave us a continuous supply of this transgenic line for experimentation. As expected, in our transgenic line only neurons in the peduncle express tdTomato, indicating that we had successfully marked ec5 neurons (Figure 1a).

**Figure 1.**
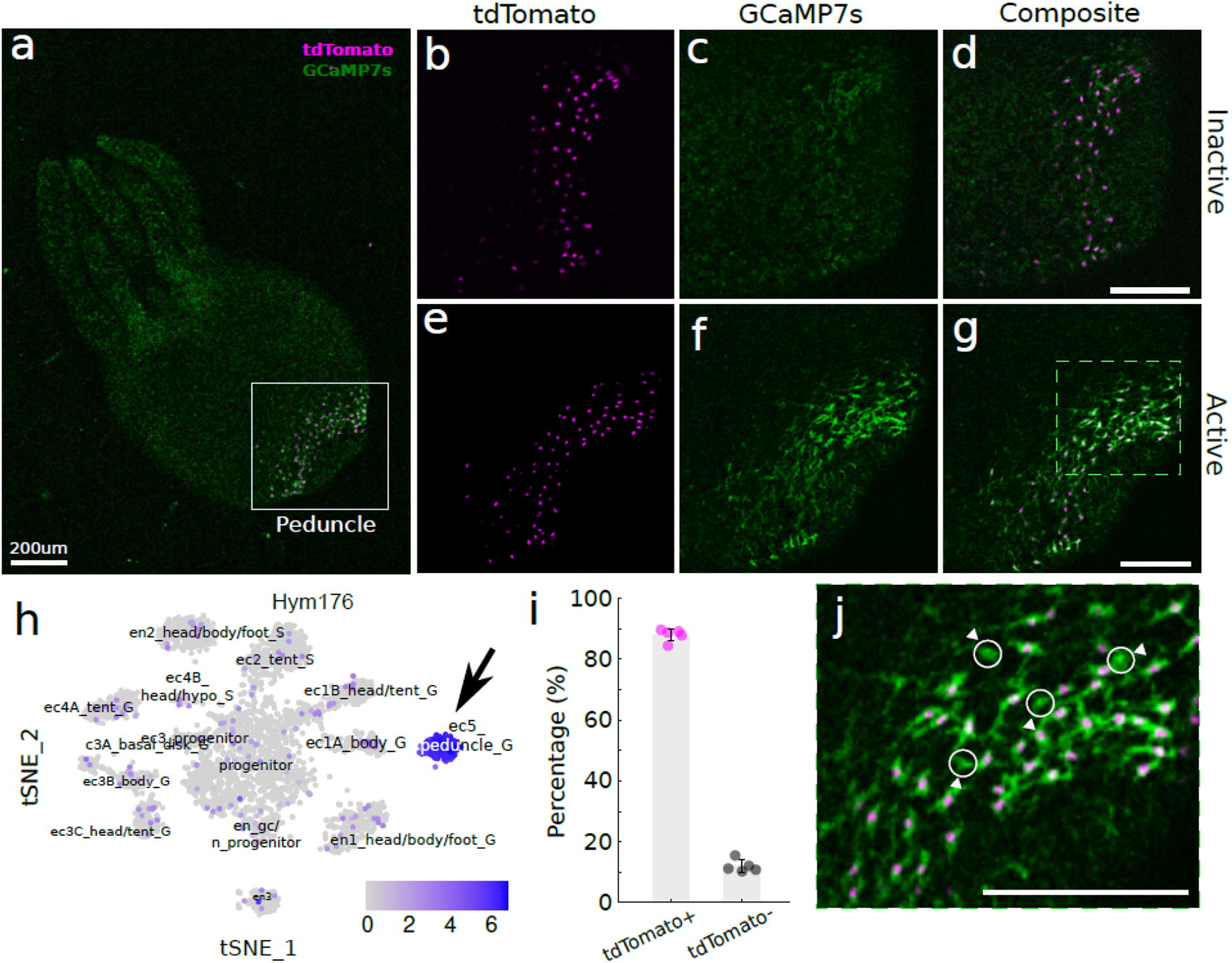
ec5 neurons are part of the Contraction Burst (CB) neural circuit. (a) Fluorescence image of a *Tg(hym176c:tdTomato,tba1c:GCAMP7s)*^cj1-in^ *Hydra* expressing nuclear-localized tdTomato in a subpopulation of neurons in the peduncle and GCaMP7s in all the neurons. Scale bar: 200um. (b-g) High magnification images of *Hydra*’s peduncle showing dual expression of tdTomato-positive neurons in magenta and GCaMP7s in green. Scale bar: 200um. (b) Nuclear tdTomato expression in the ec5 neuronal population during inactive state. (c) GCaMP7s expression in the peduncle neurons during inactive state. (d) Composite image of (b) and (c). (e) Nuclear tdTomato expression in the ec5 neuronal population during active state. (f) GCaMP7s expression in the peduncle neurons during active state. (g) Composite image of (f) and (g). (h) t-SNE representation of single neuron transcriptomes collected from *Hydra* [47]. Arrow indicates the ec5 neurons highly expressing *hym176c*. (i) The percentage of tdTomato-positive (magenta, mean = 88.02, SD = 2.08) and tdTomato-negative (black, mean = 11.98, SD = 2.08) neurons that express GCaMP7s in the peduncle. Error bars show standard deviation. (j) High magnification of (g). Circles indicate tdTomato-negative neurons in the peduncle. Scale bar: 200um.

We confirmed that GCaMP7s successfully reported calcium activity by measuring fluorescence levels during contractions. As expected, based on prior calcium imaging [12,15,32], the neuronal activity in the peduncle showed increased calcium activity during contractions (Figure 1c,f, Supplemental Video S1). We also found that the tdTomato-positive neurons were a subpopulation of the neurons that were coactive during contractions, providing further evidence that ec5 neurons are part of the CB circuit (Figure 1b-g). These ec5 neurons composed on average 88% of the neurons in the peduncle that showed increased calcium activity during contractions (n*Hydra*=5, Figure 1i). The remaining tdTomato-negative peduncle neurons (Figure 1i,j) in the CB circuit are likely the ec1A neurons that extend from the body column into the peduncle [14].

### Volumetric fluorescent imaging establishes basal levels of synchronization among neurons in the CB circuit

We used the newly created transgenic line *Tg(hym176c:tdTomato,tba1c:GCAMP7s)*^cj1-in^ to track the activity of ec5 neurons during regeneration to determine when normal circuit activity resumes. However, we first needed to establish the degree of synchrony displayed by a fully functional CB circuit in an uninjured animal. This would allow us to determine if the CB circuit has fully regenerated. To quantify the level of synchrony, we imaged the spontaneous calcium activity of tdTomato-positive ec5 neurons using SCAPE (Swept Confocally Aligned Planar Excitation) 2.0 microscopy [33]. This dual-color fast light sheet imaging technique achieves a volumetric frame rate of 1.3, which compared to the movement of *Hydra*, is fast enough to accurately track the calcium dynamics of ec5 neurons during contraction. Figure 2a,b shows multi view Maximum Intensity Projections (MIP) of the peduncle from a contracting *Tg(hym176c:tdTomato,tba1c:GCAMP7s)*^cj1-in^*Hydra* at different timepoints. The calcium activity of individual ec5 neurons was tracked (colored dots, Figure 2c) and the corresponding traces are shown in Figure 2d (Supplemental Video S1-3). We calculated the cross-correlation coefficient (CC) to measure synchrony in neural activity. Considering that the CC between the ec5 neurons (n neurons=29, n *Hydra* = 1) in an uninjured *Hydra* is 0.84 +/- 0.07 (mean +/- SEM), the CC values significantly below 0.84 in regenerating *Hydra* would indicate that the CB circuit has not yet fully recovered its function.

**Figure 2.**
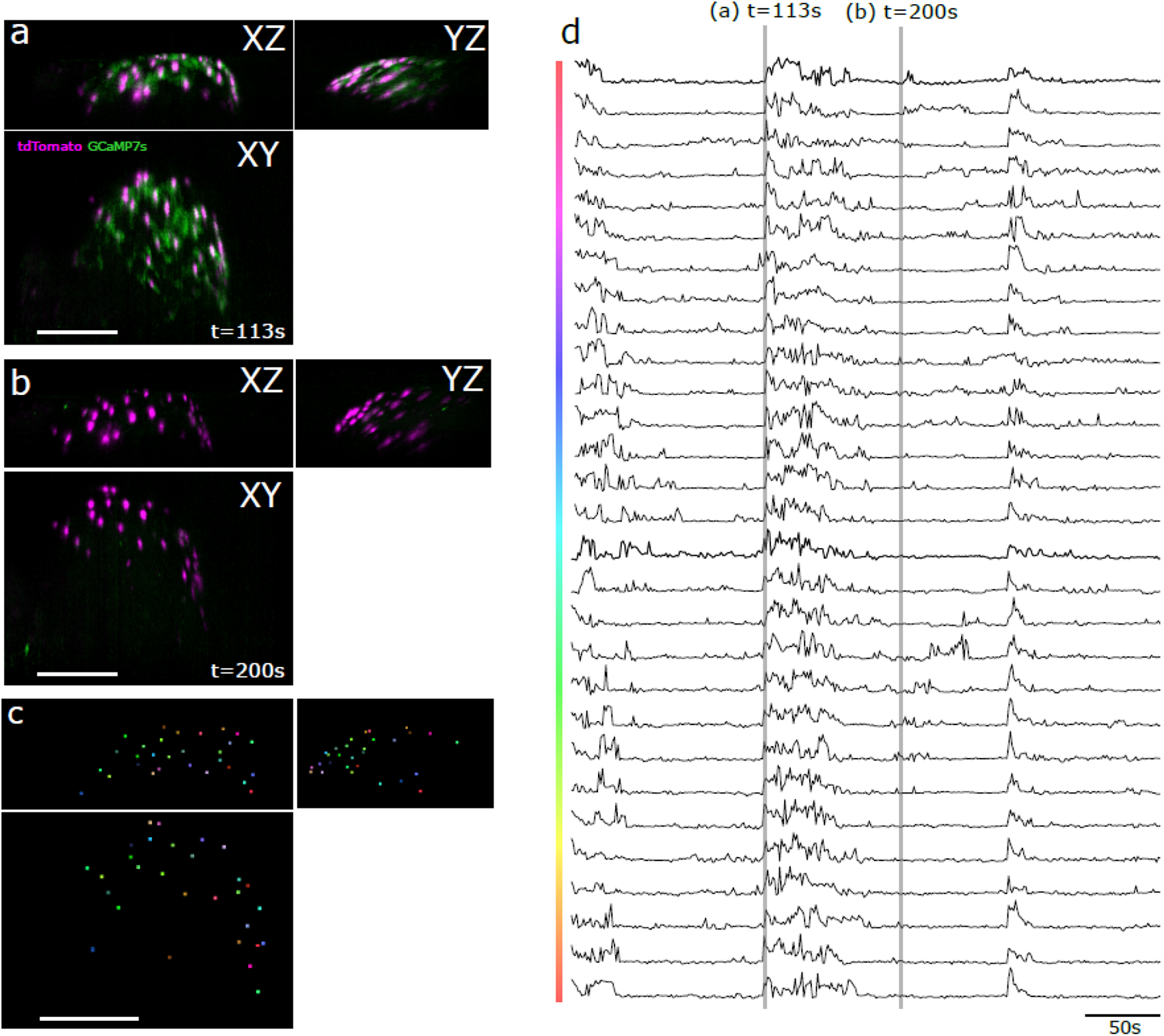
SCAPE 2.0 microscopy enables volumetric tracking of cell type specific neural activity. (a-b) Maximum intensity projections along x, y, and z axes acquired from volumetric SCAPE imaging of the peduncle of a behaving *Hydra*. Each panel shows a select time during this representative imaging session. Green shows calcium activity (GCaMP7s). Magenta shows the nuclei of ec5 cells (nuclear-localized tdTomato). Scale bar = 200 um. (c) Colored dots indicate individual neurons that are tracked over the course of recording. Scale bar = 200 um. (d) Time course of the GCaMP7s fluorescence measured in each of the labeled neurons in panels a and b. The shaded regions in gray correspond to the time points used to generate the images in panels a and b.

### c5 neuron differentiation precedes neural synchronization during regeneration

Having established a quantitative measure of synchrony in an uninjured CB circuit, we next used our new transgenic line to track the reappearance and activity of ec5 neurons during regeneration. The goal of these experiments was to determine when ec5 neurons differentiated relative to when the CB circuit resumed synchronized activity, which is indicative of a regenerated and synchronous neural circuit. We hypothesized two possible scenarios: (1) newly regenerating ec5 neurons would be functionally integrated into circuits and show a high degree of synchronization before completing terminal differentiation, or (2) ec5 neurons would express differentiation markers prior to functional integration into the CB circuit. Importantly, *hym176c* is a marker of differentiated ec5 neurons, thus the appearance of tdTomato fluorescence is a proxy for the completion of differentiation.

Since ec5 neurons are located in the peduncle [26, 37], we conducted foot regeneration experiments to track their reappearance. By bisecting the animal at the midpoint between the head and foot, we completely removed the ec5 neurons from the top half of the animal. In approximately 48 hours after this injury, the foot fully regenerates from the top half (Figure 3a). During this time, new neurons are produced from the interstitial stem cells that reside among the ectodermal epithelial cells in the body column. Over the course of regeneration we tracked the reappearance of ec5 neurons (using tdTomato expression) and the activity of the CB circuit in the peduncle (using GCaMP7s fluorescence) by taking 20 minute recordings every four hours post amputation (hpa) (n = 5 animals) to monitor circuit reformation.

**Figure 3.**
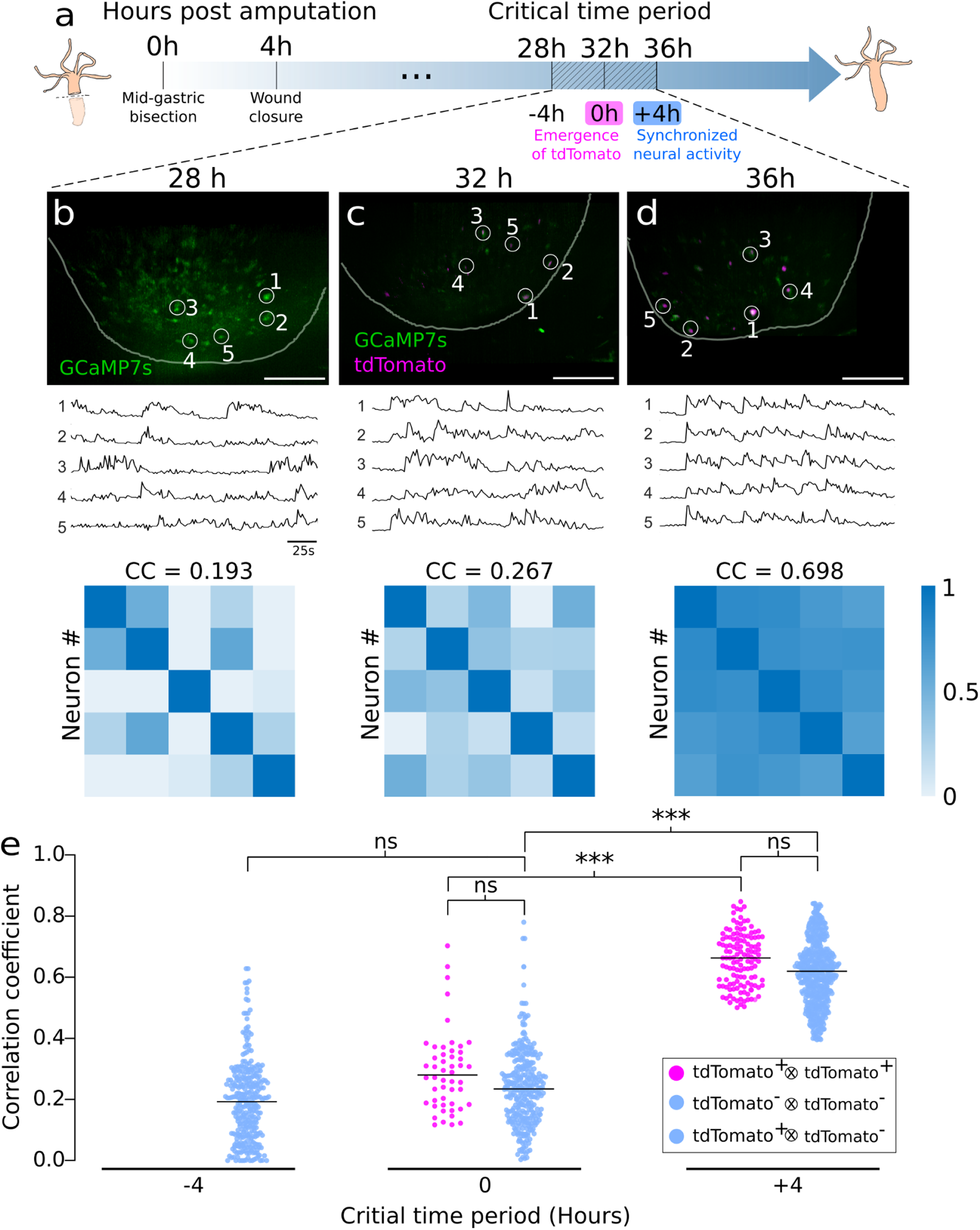
ec5 neuron differentiation precedes synchronization in neural activity during peduncle regeneration. (a) Schematic representation of foot regeneration timeline after a mid-gastric bisection. (b-d) (Top row) Representative composite fluorescence image of nuclear tdTomato (magenta) and GCaMP7s (green) from the same *Hydra* at indicated time points, scale bar = 100 μm. (Middle row) GCaMP7s traces extracted from the circled neurons at time points corresponding to the top row. Numbered circles indicate the neurons of which spontaneous GCaMP7s activities are plotted. (Bottom row) Cross-correlation matrix of the circled neurons as an indicator for synchrony at time points corresponding to the top and middle row with average correlation coefficient (CC). (e) Dots represent correlation levels between two arbitrary neurons’ activity. Correlation between two tdTomato-positive neurons is labeled pink. Correlation between two tdTomato-negative neurons, or between one tdTomato-positive and one tdTomato-negative neuron is labeled blue. (ns = not significant, * p ≤ 0.05, ** p ≤ 0.01, *** p ≤ 0.001, Student’s t test with bonferroni correction).

In a representative *Hydra* shown in Figure 3, we randomly selected 5 neurons from *Tg(hym176c:tdTomato,tba1c:GCAMP7s)*^cj1-in^ *Hydra* to evaluate synchrony. We observed unsynchronized neural activity with no tdTomato-positive neurons (Figure 3b, Supplemental Video S4) from 0-28 hpa. At 32 hpa, the tdTomato-positive neurons began to appear, but these neurons exhibited a low level of synchrony (Figure 3c, Supplemental Video S5). At the 36 hpa mark, we observed an increase in the number of tdTomato-positive neurons along with a large increase in synchronization (Figure 3d, Supplemental Video S6). Due to the varying timing of the appearance of tdTomato-positive neurons across multiple animals, we defined the time point when the tdTomato-positive (ec5) neurons first reappeared as t = 0hr (n *Hydra* = 5). With this alignment, we found a critical window of four hours that separates the first detection of tdTomato-positive cells and the synchronization of the CB circuit, named the “critical time period” (Figure 3a). At t = -4hr, the activity of the neurons in the regenerating foot were not synchronized (Figure 3e, correlation coefficient = 0.172+/-0.07 (mean +/- SEM)). At t = 0hr, the synchrony of the neurons was low (correlation coefficient = 0.239+/-0.07) even though tdTomato-positive neurons appeared. The level of synchrony increased (correlation coefficient = 0.609+/-0.03) at t = +4hr where we also observed an increase in the number of tdTomato-positive neurons. However, they were less synchronized compared to the neurons in uninjured animals (0.84 +/- 0.07 (mean +/- SEM)). Together, these data support the scenario in which ec5 neurons fully differentiate before functional integration into the CB circuit.

## Conclusion

Although specification of neurons and assembly of neural circuits during development is relatively well studied, these processes are not as well understood during regeneration. Several research organisms, including zebrafish (*Danio rerio)*, axolotl (*Ambystoma mexicanum*), xenopus tadpoles, and planarians (*Schmidtea mediterranea)*, can regenerate large portions of their bodies including innervation and the restoration of behavior [37,43,44,45,46]. These animals have provided several interesting insights, including the ability of zebrafish [8] and xenopus tadpoles [42] to restore neural circuit activity and behavior before regeneration is complete (cite). *Hydra* has a unique combination of advantages as compared to existing neuronal regeneration models which allows us to image the activity of the entire nervous system at single cell resolution. In this study, we build new tools and leverage volumetric imaging to examine the interplay between neuronal differentiation and the resumption of neural circuit activity during *Hydra* nervous system regeneration. We find that the ec5 neurons reach their terminal cell-fate before they functionally integrate into the *Hydra* CB circuit. These data suggest that the cues from surrounding cells direct differentiation of stem cells into the appropriate neuron subtypes rather than existing circuit activity directing these fates. Further work should be done to identify these injury-induced differentiation signals.

This work also raises the question of whether cell fate determination prior to functional recovery is true in other circuits in *Hydra*. The platform and approach developed here provide a powerful tool for answering this and other questions in *Hydra*. In particular a major accomplishment of this work is the first demonstration of dual reporter expression in *Hydra* neurons that allows for the combined measurement of neural activity and gene expression in a live animal during regeneration. Combining this reporter system with high-speed volumetric imaging brings *Hydra* into the small but growing group of neuroscience research organisms [37,43,44] in which we can perform functional volumetric imaging with cell-type specificity. To apply this method to other circuits in *Hydra*, future work should identify quantitative measures of neural activity that indicate functional recovery in each circuit. The ability to image neuronal activity throughout the animal during behavior along with cell type specific labeling will allow one to collect the types of data needed to identify normal neural circuit activity. Overall these transgenic and imaging tools combined with *Hydra’s* unique regenerative ability differentiates it with respect from other research organisms as a model for studying complete neural circuit regeneration.

## Methods

### Generation of *Tg(hym176c:tdTomato,tba1c:GCAMP7s)*^cj1-in^ transgenic strain

*Hydra* transgenic line *Tg(hym176c:tdTomato,tba1c:GCAMP7s)*^cj1-in^ was created by microinjecting a single plasmid containing two promoters and two transgenes. Nuclear tdTomato was driven by a 2022 bp section of the *hym176c* regulatory region, which should be specifically expressed in ec5 neurons found in the peduncle (Fig 1H, 26), and GCaMP7s was driven by a 1901 bp section of the *tba1c* regulatory region, which was validated in Primack et al. (2023) to be a pan-neuronal promoter. The plasmid injection solution was injected into *Hydra vulgaris* AEP 1-cell stage embryos using an Eppendorf FemtoJet 4x and Eppendorf InjectMan NI 2 microinjector (Eppendorf; Hamburg, Germany) under a Leica M165 C stereo microscope (Leica Microscopes, Inc; Buffalo Grove, Il).

Hatchlings were screened to select tdTomato-positive polyps. Continuous asexual reproduction cycles of hatchlings with mosaic transgenic tissue yielded transgenic animals with uniform gene expression. The DNA plasmid was designed by the Robinson Lab (Rice University) and the transgenic strain was developed by Celina Juliano’s Laboratory (University of California, Davis) following an established protocol [14].

### *Hydra* strain maintenance

All animals were maintained using standard procedures (Lenhoff & Brown, 1970). All experiments were performed using the transgenic *Hydra* line *Tg(hym176c:tdTomato,tba1c:GCAMP7s)*^cj1-in^. *Hydra* polyps were cultured at standard conditions, incubated at 18°C with *Hydra* media under 12hr:12hr light:dark light cycles. *Hydra* media was made with 1000X dilution of 1.0M CaCl_2_, 0.1M MgCl_2_, 0.03M KNO_3_, 0.5M NaHCO_3_, 0.08M MgSO_4_. Polyps were fed three times per week with freshly hatched *Artemia nauplii* (Brine Shrimp Direct) and cleaned 6 hours post feeding with *Hydra* media. The animals were starved 24h hours prior to surgical resections.

### Animal resections

*Hydra* were placed in a petri dish filled with *Hydra* media prior to the incision. Animal resections were performed using a scalpel, making a single incision across the center of the body, removing the aboral end and keeping the oral end to track the foot regeneration. Resectioned *Hydra* were kept at 18°C for 4 hours to allow wound closure before subjecting them to imaging procedures.

### Imaging configuration

Imaging was performed by placing a resected *Hydra* between two glass coverslips separated by a 100 um spacer. Dual-color volumetric imaging was performed in 20 minute sessions at a 4 hour interval for 44 hours post amputation (hpa) at 100 fps (1.3 VPS) using Swept Confocally Aligned Planar Excitation (SCAPE) 2.0 microscopy. The system was built following the design and configuration from Voleti et al. with assistance and support from Elizabeth Hillman’s laboratory at Columbia University [33]. The configuration consisted of an 20X Olympus (XLUMPLFLN 20XW 20x/1.00NA) as the primary objective lens, (for specimen illumination and light collection), followed by a Nikon 20x/0.75NA and Nikon 10x/0.45NA as the second and third objective lenses according to the SCAPE system nomenclature. The system used in all experiments had an effective detection NA of 0.23 and used Andor Zyla 4.2+ as the detector for imaging sessions. The microscope system provided oblique light sheet illumination across the field of view (800um × 350um × 100m). Oblique illumination was achieved by enabling the light sheet to enter the back aperture of the primary objective lens with an offset of 7 mm from the center of the objective. Coherent Obis LX 488 nm and 561 nm lasers were used as excitation laser sources for green (GCaMP7s) and red (tdTomato) channels respectively at an output power of 5mW/mm^2^. Excitation and emission filters used for all dual color imaging experiments are listed in Supplementary Table 1. Epifluorescence microscopy was used for whole-animal imaging (Figure 1).

### Image processing and cell tracking

For injured *Hydra*, both channels were acquired and registered to one another and exported to 16 bit tiff format using a custom MATLAB GUI provided by Elizabeth Hillman’s laboratory at Columbia University. Imaris software and 3D Viewer plugin from Fiji [49] were used to visualize the channel-merged image sequence in 3D. Then, maximum intensity projections from recordings were imported to Fiji and contrast was adjusted. Particle-tracking algorithm TrackMate v6.0.2 and ManualTracking [48] plugin from Fiji were used to track single cell activity. Prior to tdTomato expression, single cell GCaMP7s time courses were manually annotated using frame to frame analysis (ManualTracking) from Fiji. Posterior to tdTomato expression, GCaMP7s traces corresponding to tdTomato-negative neurons continued to be manually annotated using ManualTracking while GCaMP7s traces from tdTomato-positive neurons were automatically tracked using TrackMatev6.0.2 (LoG detector sigma:15-20; Simple LAP tracker) followed by manual corrections.

For uninjured *Hydra* both channels were acquired and registered to one another and exported to 16 bit tiff format using a custom MATLAB GUI provided by Elizabeth Hillman’s laboratory at Columbia University. Imaris software was used to visualize the channel-merged image sequence in 3D. Autoregressive motion with MaxGapSize=3 was used for particle tracking. Cells that fall outside the FOV for more than 50% of the recorded time were not included in the tracking process.

### Statistical analysis of neural activity

For all animals (n = 5) single cell calcium activity traces from all regeneration time points (8 - 44 hpa, Supplemental figure 1) were analyzed using MATLAB (Figure 3). Relative GCaMP7s intensity was normalized using the minimum and maximum pixel intensity. To evaluate the level of synchrony in the neurons of interest, all obtained traces were compared to each other by measuring the linear dependence between two arbitrary cells’ activity and assigning a Pearson correlation coefficient (calculated with the MATLAB function corrcoef). The linear relationship between two arbitrary time series was performed on 15 min time series. From a scale of 0 to 1, higher value indicates higher correlation. The obtained correlation coefficients were used to build a correlation matrix for every regeneration time point. To compare the difference of correlation coefficients within and between the regeneration time points we performed an unpaired Student’s t test with bonferroni corrections.

## Supplemental figures

**Supplemental Figure 1.**
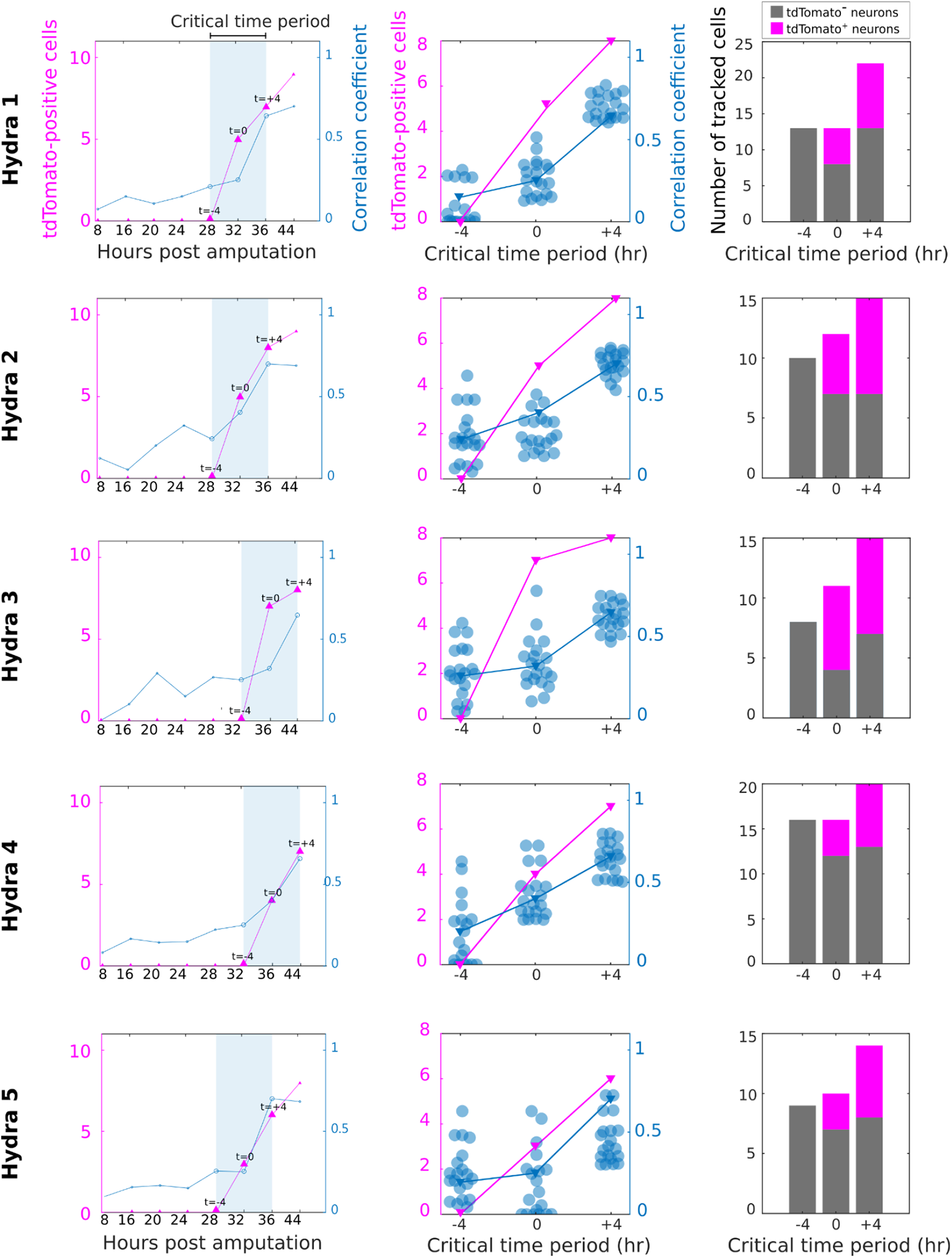
Cell count and synchrony in neural activity for n = 5 *Hydra*. (Left column) The number of TdTomato-positive neurons in magenta, and the average correlation coefficient (CC) tracked over the course of 8 - 44 hpa. The blue shaded region indicates the critical time period. (Middle column) The number of tdTomato-positive neurons in magenta and the CC values in blue with average shown in inverted triangle during the critical time period. (Right panel) The number of tracked cells during the critical time period.

**Supplemental Video 1**. Single cell activity of an uninjured *Hydra’s* foot. Playback speed 1X.

**Supplemental Video 2**. Single cell tracking of an uninjured *Hydra’s* foot. Playback speed 1X.

**Supplemental Video 3**. Single cell activity traces of an uninjured *Hydra’s* foot. Playback speed 1X. 2 cells that were outside the FOV for more than >67% of the recorded time were not included in the tracking process.

**Supplemental Video 4**. Single cell activity of an injured *Hydra* at the critical time point “t=-4hr” showing unsynchronized activity of GCaMP7s-positive cells and absence of ec5 neurons. Playback speed 1X.

**Supplemental Video 5**. Single cell activity of a regenerating *Hydra* at the critical time point “t=0hr” showing unsynchronized activity of GCaMP7s-positive cells and emergence of ec5 neurons. Playback speed 1X.

**Supplemental Video 6**. Single cell activity of a regenerating *Hydra* at the critical time point “t=+4hr” showing synchronized activity of GCaMP7s-positive cells and ec5 neurons. Playback speed 1X.

**Supplementary Table 1.** Optic filters used in regeneration experiments

| Filter type | Description | Vendor |
| --- | --- | --- |
| Excitation/emission dichroic | 500 nm long pass | FF495-Di03-25x36<br>(Semrock) |
| Emission | 495 nm long-pass dichroic | FF01-496/LP-25<br>(Semrock) |
| Image splitter dichroic | 560 nm long-pass dichroic | FF560-FDi01-25x36<br>(Semrock) |
| Green emission filter | 525 +/- 50 nm bandpass | FF01-525/50-25<br>(Semrock) |
| Red emission filter | 618 +/- 50 nm bandpass | FF01-618/50-25<br>(Semrock) |
| Laser combining dichroic | 556 nm short-pass dichroic | FF556-SDi01-25x36<br>(Semrock) |
| Dichroic Beam Splitter | 488/561 nm dichroic beamsplitter | Di01-R488/561-25x36<br>(Semrock) |

## References

1. Peter W. Reddien, “The Cellular and Molecular Basis for Planarian Regeneration,” Cell 175, no. 2 (October 2018): 327–45, https://doi.org/10.1016/j.cell.2018.09.021.

2. Daniel Lobo, Wendy S. Beane, and Michael Levin, “Modeling Planarian Regeneration: A Primer for Reverse-Engineering the Worm,” ed. Takashi Gojobori, PLoS Computational Biology 8, no. 4 (April 26, 2012): e1002481, https://doi.org/10.1371/journal.pcbi.1002481.

3. Michael P Sarras Jr, “Hydra as a Unique Model for the Study of Regenerative Mechanisms in Metazoans,” MOJ Anatomy & Physiology 6, no. 5 (September 16, 2019), https://doi.org/10.15406/mojap.2019.06.00264.

4. Puli Chandramouli Reddy, Akhila Gungi, and Manu Unni, “Cellular and Molecular Mechanisms of Hydra Regeneration,” in Evo-Devo: Non-Model Species in Cell and Developmental Biology, ed. Waclaw Tworzydlo and Szczepan M. Bilinski, vol. 68, Results and Problems in Cell Differentiation (Cham: Springer International Publishing, 2019), 259–90, https://doi.org/10.1007/978-3-030-23459-1_12.

5. Matthias C. Vogg, Brigitte Galliot, and Charisios D. Tsiairis, “Model Systems for Regeneration: Hydra,” Development 146, no. 21 (November 1, 2019): dev177212, https://doi.org/10.1242/dev.177212.

6. Warren A. Vieira, Kaylee M. Wells, and Catherine D. McCusker, “Advancements to the Axolotl Model for Regeneration and Aging,” Gerontology 66, no. 3 (2020): 212–22, https://doi.org/10.1159/000504294.

7. Kaylee M Wells et al., “Neural Control of Growth and Size in the Axolotl Limb Regenerate,” eLife 10 (November 15, 2021): e68584, https://doi.org/10.7554/eLife.68584.

8. Celia Vandestadt et al., “RNA-Induced Inflammation and Migration of Precursor Neurons Initiates Neuronal Circuit Regeneration in Zebrafish,” Developmental Cell 56, no. 16 (August 2021): 2364-2380.e8, https://doi.org/10.1016/j.devcel.2021.07.021.

9. Giorgia Beffagna, “Zebrafish as a Smart Model to Understand Regeneration After Heart Injury: How Fish Could Help Humans,” Frontiers in Cardiovascular Medicine 6 (August 6, 2019): 107, https://doi.org/10.3389/fcvm.2019.00107.

10. Ziad Julier et al., “Promoting Tissue Regeneration by Modulating the Immune System,” Acta Biomaterialia 53 (April 2017): 13–28, https://doi.org/10.1016/j.actbio.2017.01.056.

11. Bai Lu, Kuan Hong Wang, and Akinao Nose, “Molecular Mechanisms Underlying Neural Circuit Formation,” Current Opinion in Neurobiology 19, no. 2 (April 2009): 162–67, https://doi.org/10.1016/j.conb.2009.04.004.

12. Matthias C. Vogg, Brigitte Galliot, and Charisios D. Tsiairis, “Model Systems for Regeneration: Hydra,” Development 146, no. 21 (November 1, 2019): dev177212, https://doi.org/10.1242/dev.177212.

13. S. E. Wilson-Sanders, “Invertebrate Models for Biomedical Research, Testing, and Education,” ILAR Journal 52, no. 2 (January 1, 2011): 126–52, https://doi.org/10.1093/ilar.52.2.126.

14. Jörg Wittlieb et al., “Transgenic Hydra Allow in Vivo Tracking of Individual Stem Cells during Morphogenesis,” Proceedings of the National Academy of Sciences 103, no. 16 (April 18, 2006): 6208–11, https://doi.org/10.1073/pnas.0510163103.

15. Jarrod A. Chapman et al., “The Dynamic Genome of Hydra,” Nature 464, no. 7288 (March 2010): 592–96, https://doi.org/10.1038/nature08830.

16. Thomas C.G. Bosch et al., “Back to the Basics: Cnidarians Start to Fire,” Trends in Neurosciences 40, no. 2 (February 2017): 92–105, https://doi.org/10.1016/j.tins.2016.11.005.

17. John R. Szymanski and Rafael Yuste, “Mapping the Whole-Body Muscle Activity of Hydra Vulgaris,” Current Biology 29, no. 11 (June 2019): 1807-1817.e3, https://doi.org/10.1016/j.cub.2019.05.012.

18. John F. Dunne et al., “A Subset of Cells in the Nerve Net of Hydra Oligactis Defined by a Monoclonal Antibody: Its Arrangement and Development,” Developmental Biology 109, no. 1 (May 1985): 41–53, https://doi.org/10.1016/0012-1606(85)90344-6.

19. Christophe Dupre and Rafael Yuste, “Non-Overlapping Neural Networks in Hydra Vulgaris,” Current Biology 27, no. 8 (April 2017): 1085–97, https://doi.org/10.1016/j.cub.2017.02.049.

20. Nathan De Fruyt et al., “The Role of Neuropeptides in Learning: Insights from C. Elegans,” The International Journal of Biochemistry & Cell Biology 125 (August 2020): 105801, https://doi.org/10.1016/j.biocel.2020.105801.

21. Seth R. Taylor et al., “Molecular Topography of an Entire Nervous System,” Cell 184, no. 16 (August 2021): 4329-4347.e23, https://doi.org/10.1016/j.cell.2021.06.023.

22. Yukihiko Noro et al., “Regionalized Nervous System in Hydra and the Mechanism of Its Development,” Gene Expression Patterns 31 (January 2019): 42–59, https://doi.org/10.1016/j.gep.2019.01.003.

23. Yukihiko Noro et al., “A Single Neuron Subset Governs a Single Coactive Neuron Circuit in Hydra Vulgaris, Representing a Possible Ancestral Feature of Neural Evolution,” Scientific Reports 11, no. 1 (May 24, 2021): 10828, https://doi.org/10.1038/s41598-021-89325-x.

24. Toshio Takahashi, “Hym-176,” in Handbook of Hormones (Elsevier, 2016), 485–86, https://doi.org/10.1016/B978-0-12-801028-0.00090-8.

25. GN Hansen, Michael Williamson, and Cornelis J.P. Grimmelikhuijzen, “Two-Color Double-Labeling in Situ Hybridization of Whole-Mount Hydra Using RNA Probes for Five Different Hydra Neuropeptide Preprohormones: Evidence for Colocalization,” Cell and Tissue Research 301, no. 2 (July 19, 2000): 245–53, https://doi.org/10.1007/s004410000240.

26. Stefan Siebert et al., “Stem Cell Differentiation Trajectories in Hydra Resolved at Single-Cell Resolution,” Science 365, no. 6451 (July 26, 2019): eaav9314, https://doi.org/10.1126/science.aav9314.

27. Ulf Knoblich et al., “Neuronal Cell-Subtype Specificity of Neural Synchronization in Mouse Primary Visual Cortex,” Nature Communications 10, no. 1 (June 10, 2019): 2533, https://doi.org/10.1038/s41467-019-10498-1.

28. Romain Brette, “Computing with Neural Synchrony,” ed. Olaf Sporns, PLoS Computational Biology 8, no. 6 (June 14, 2012): e1002561, https://doi.org/10.1371/journal.pcbi.1002561.

29. Peter Uhlhaas, “Neural Synchrony in Cortical Networks: History, Concept and Current Status,” Frontiers in Integrative Neuroscience 3 (2009), https://doi.org/10.3389/neuro.07.017.2009.

30. Hod Dana et al., “High-Performance Calcium Sensors for Imaging Activity in Neuronal Populations and Microcompartments,” Nature Methods 16, no. 7 (July 2019): 649–57, https://doi.org/10.1038/s41592-019-0435-6.

31. Celina E. Juliano, Haifan Lin, and Robert E. Steele, “Generation of Transgenic Hydra by Embryo Microinjection,” Journal of Visualized Experiments, no. 91 (September 11, 2014): 51888, https://doi.org/10.3791/51888.

32. Krishna N Badhiwala et al., “Multiple Neuronal Networks Coordinate Hydra Mechanosensory Behavior,” eLife 10 (July 30, 2021): e64108, https://doi.org/10.7554/eLife.64108.

33. Venkatakaushik Voleti et al., “Real-Time Volumetric Microscopy of in Vivo Dynamics and Large-Scale Samples with SCAPE 2.0,” Nature Methods 16, no. 10 (October 2019): 1054–62, https://doi.org/10.1038/s41592-019-0579-4.

34. Kiyokazu Agata, Yumi Saito, and Elizabeth Nakajima, “Unifying Principles of Regeneration I: Epimorphosis versus Morphallaxis,” Development, Growth & Differentiation 49, no. 2 (February 2007): 73–78, https://doi.org/10.1111/j.1440-169X.2007.00919.x.

35. Andrey Elchaninov, Gennady Sukhikh, and Timur Fatkhudinov, “Evolution of Regeneration in Animals: A Tangled Story,” Frontiers in Ecology and Evolution 9 (March 5, 2021): 621686, https://doi.org/10.3389/fevo.2021.621686.

36. Alejandro Sánchez Alvarado, “Stem Cells in Animal Models of Regeneration,” StemBook, 2008, https://doi.org/10.3824/stembook.1.32.1.

37. Alessandro Zambusi and Jovica Ninkovic, “Regeneration of the Central Nervous System-Principles from Brain Regeneration in Adult Zebrafish,” World Journal of Stem Cells 12, no. 1 (January 26, 2020): 8–24, https://doi.org/10.4252/wjsc.v12.i1.8.

38. Sukla Ghosh and Subhra Prakash Hui, “Regeneration of Zebrafish CNS: Adult Neurogenesis,” Neural Plasticity 2016 (2016): 1–21, https://doi.org/10.1155/2016/5815439.

39. Andrew R Gehrke and Mansi Srivastava, “Neoblasts and the Evolution of Whole-Body Regeneration,” Current Opinion in Genetics & Development 40 (October 2016): 131–37, https://doi.org/10.1016/j.gde.2016.07.009.

40. Christian P. Petersen, “Regeneration: Organizing the Blastema in Planarians,” Current Biology 27, no. 5 (March 2017): R181–83, https://doi.org/10.1016/j.cub.2017.01.057.

41. Athina Keramidioti et al., “A new look at the architecture and dynamics of the Hydra nerve net” (2023) bioRxiv doi: https://doi.org/10.1101/2023.02.22.529525.

42. Anneke Dixie Kakebeen, et al., “Chromatin accessibility dynamics and single cell RNA-Seq reveal new regulators of regeneration in neural progenitors,” eLife 9:e52648, https://doi.org/10.7554/eLife.52648.

43. Lee-Liu D, et al., “The African clawed frog Xenopus laevis: A model organism to study regeneration of the central nervous system,” Neurosci Lett. (2017) Jun 23;652:82–93. doi: 10.1016/j.neulet.2016.09.054. Epub 2016 Sep 29. PMID: 27693567.

44. Ross KG, et al., RM. “Nervous system development and regeneration in freshwater planarians,” Wiley Interdiscip Rev Dev Biol. 2017 May;6(3). doi: 10.1002/wdev.266. Epub 2017 Mar 22. PMID: 28326682.

45. Wei X, et al. “Single-cell Stereo-seq reveals induced progenitor cells involved in axolotl brain regeneration,” Science 2022 Sep 2;377(6610):eabp9444. doi: 10.1126/science.abp9444. Epub 2022 Sep 2. PMID: 36048929.

46. Lust K, et al., “Single-cell analyses of axolotl telencephalon organization, neurogenesis, and regeneration,” Science. 2022 Sep 2;377(6610):eabp9262. doi: 10.1126/science.abp9262. Epub 2022 Sep 2. PMID: 36048956.

47. Abby S. Primack, et al., “Differentiation trajectories of the Hydra nervous system reveal transcriptional regulators of neuronal fate” (2023) bioRxiv doi: https://doi.org/10.1101/2023.03.15.531610.

48. Tinevez, J.-Y.,et al., (2017). TrackMate: An open and extensible platform for single-particle tracking. Methods, 115, 80–90. doi:10.1016/j.ymeth.2016.09.016.

49. Schindelin, J.,et al., (2012). Fiji: an open-source platform for biological-image analysis. Nature Methods, 9(7), 676–682. doi:10.1038/nmeth.2019

